# EFFECT OF A LACTOBACILLUS POSTBIOTIC ON DIGESTIVE HEALTH IN DOGS RAISED IN CONDITIONS AT RISK OF *GIARDIA DUODENALIS* INFECTION

**DOI:** 10.64898/2026.11.17.767253

**Authors:** Erik Eckhardt, Sergio Andreu-Sánchez, Marco van den Berg, Anne-Sophie Boucard, Balint Tamasi, Brian Klein, Elena Pagani, Dany Espuche, Aurélie Chevillot, Maxime Kurtz, Benedicte Lagofun, Pierre Deshuillers, Delphine Le Roux, Noelle Cochet-Faivre, Bruno Polack, Mohamed Mammeri

## Abstract

2.

*Giardia duodenalis* infection is associated with diarrhea and dysbiosis in dogs, while metronidazole treatment can further disrupt the microbiota. We evaluated adjunctive LB fermentate HF (LBHF), a Lactobacillus-derived postbiotic, in a randomized, blinded, placebo-controlled study of 30 naturally infected dogs (Terriers). Animals received metronidazole (50 mg/kg/day) with LBHF (50 mg/kg/day) or microcrystalline cellulose placebo (70 mg/kg/day) for 21 days. Clinical outcomes, fecal biomarkers, canine dysbiosis index, and cyst shedding were monitored over 45 days. Fecal shotgun metagenomics was performed at baseline and day 45.

After baseline covariate adjustment, LBHF was associated with significantly lower *Giardia* burdens during treatment, particularly at days 10 and 20, improved fecal consistency, and reduced odor. No significant effects were found for fecal calprotectin, secretory IgA, blood biomarkers of digestive function, or canine dysbiosis index. Overall microbiome turnover was similar between groups, but multivariate analyses identified distinct treatment-associated community trajectories. LBHF was associated with increased Bacteroidota representation and functional capacity for plant polysaccharide degradation, alongside depletion of gene sets involved in flagellar assembly, quorum sensing, biofilm formation, and epithelial-cell signaling. Baseline microbiome composition also predicted *Giardia* burden, highlighting an association between parasite load and bacterial community structure.

Adjunctive LBHF supplementation was associated with faster resolution of canine giardiasis, improved fecal quality, and coordinated microbiome remodeling. Further mechanistic studies are needed to establish whether the observed microbiome functional changes contribute to parasite clearance and recovery of intestinal health.

**Highlights:**

- A Lactobacillus postbiotic was linked to reduced *Giardia* cyst shedding in infected dogs
- The postbiotic also improved fecal consistency and reduced fecal odor.
- The postbiotic was linked to greater potential for plant-polysaccharide degradation.
- Baseline microbiome composition was associated with *Giardia* burden

## 5. Introduction

Giardiosis, caused by the intestinal protozoan parasite *Giardia duodenalis* (syn. *G. intestinalis*, *G. lamblia*), is one of the most prevalent gastrointestinal parasitic infections in dogs worldwide, with reported prevalence rates ranging from 5% to over 50% depending on the population studied and the diagnostic method employed (Šlapeta et al., 2015; Sui et al., 2022). Young animals, particularly puppies raised in kennels or shelters, are disproportionately affected (Agresti et al., 2022; Fontanarrosa et al., 2006; Kurnosova et al., 2024), and clinical manifestations range from asymptomatic cyst shedding to acute or chronic diarrhea, malabsorption, weight loss, and growth retardation. The infection is caused by ingestion of environmentally resistant cysts that excyst in the small intestine, where trophozoites colonize the brush border and disrupt epithelial barrier function, nutrient absorption, and local immune responses (Fekete et al., 2020).

Beyond its direct pathological effects, *G. duodenalis* infection in dogs has been consistently associated with dysbiosis, which is characterized by a reduction in the abundance of beneficial commensal species and an increase in pathogenic and pro-inflammatory taxa (Berry et al., 2020; Boucard et al., 2021; Šlapeta et al., 2015; Sui et al., 2022). A landmark study conducted in Australia on 40 dogs showed that *Giardia* induced markedly more pronounced microbiota alterations than other parasites such as hookworms (*Ancylostoma*) or coccidia (*Cystoisospora*) (Šlapeta et al., 2015), a finding subsequently corroborated in a larger cohort of 258 dogs (Berry et al., 2020). A longitudinal study conducted in a French Beagle breeding facility further demonstrated that puppies with high *Giardia* cyst excretion displayed microbiota changes, characterized by increased numbers of facultative anaerobic, mucus-degrading and pro-inflammatory species, and depletion of *Lactobacillus* compared with low-shedding animals (Boucard et al., 2021). Importantly, dysbiosis may persist after parasitological treatment and has been proposed as a driver of chronic diarrhea observed in some individuals (Ciuca et al., 2021; Fekete et al., 2020). *G. duodenalis* thus represents a particularly instructive model for studying parasite-microbiome interactions and for evaluating microbiota-targeted therapeutics in dogs.

Current first-line treatment for canine giardiasis relies primarily on antiparasitic agents, most notably metronidazole (50 mg/kg/day), which achieves variable parasitological clearance rates and is frequently used in combination with other antiparasitic drugs such as fenbendazole (Ciuca et al., 2021). However, pharmacological treatment carries an important limitation: metronidazole is itself a broad-spectrum antimicrobial that substantially disrupts the intestinal microbiota, reducing microbial diversity and selectively depleting anaerobic bacteria (Marshall-Jones et al., 2024; Pilla et al., 2020). Thus, metronidazole-based therapy may paradoxically exacerbate dysbiosis, potentially contributing to post-treatment clinical relapses and delayed microbiota recovery.

Several nutritional strategies have been investigated to mitigate intestinal dysbiosis in dogs, including dietary fiber supplementation, probiotics, and more recently, postbiotics (Belà et al., 2024; Bonel-Ayuso et al., 2025; Schmitz C Suchodolski, 2016; Xia et al., 2024). According to the 2021 consensus definition of the International Scientific Association for Probiotics and Prebiotics (ISAPP), a postbiotic is defined as a "preparation of inanimate microorganisms and/or their components that confers a health benefit on the host" (Salminen et al., 2021). Unlike live probiotics, postbiotics are inanimate microbial preparations that are more stable than live bacteria and pose a reduced risk upon systemic translocation compared to live microbials. Postbiotics promote intestinal health by reinforcing epithelial barrier function and modulating mucosal immune responses, with demonstrated effects on gut microbiota composition in multiple species. In dogs, a *Lactobacillus*-based fermentation postbiotic (LB fermentate HF (“LBHF”), dsm-firmenich Houdan France) supplemented at 4 mg/kg/day for 5 weeks improved fecal consistency, significantly reduced oxidative stress, and increased the abundance of beneficial bacteria including *Bifidobacterium* in adult Beagles (Koziol et al., 2023). In pigs orally challenged with *Escherichia coli* F18+, the same product reduced post-weaning diarrhea and improved intestinal histomorphology (Duarte et al., 2024; Xu et al., 2022), corroborating ex-vivo observations of gut barrier strengthening (Cercamondi et al., 2025). In murine models, the same postbiotic modulated the gut microbiome and reduced *Citrobacter*-induced colitis (Warda et al., 2020).

The present study was designed to evaluate the association between LBHF supplementation, administered adjunctively to conventional metronidazole treatment, and the clinical and microbiological course of *G. duodenalis* infection in naturally infected puppies raised in conditions at high risk of parasitic exposure. The dual rationale for this intervention was: (i) to counteract microbiota disruption induced by metronidazole, and (ii) to restore the dysbiosis associated with *Giardia* infection itself. The primary objective of the study was to assess the association between LBHF plus metronidazole (*versus* metronidazole alone) and *Giardia* cyst clearance kinetics in canine puppies with naturally occurring *G. duodenalis* infection. Secondary objectives included evaluation of the effect of LBHF plus metronidazole (*versus* metronidazole alone) on fecal scores (BSS and odor score), fecal inflammatory biomarkers (calprotectin, secretory IgA), the canine dysbiosis index, blood biomarkers of digestive function, serum cortisol concentration, and gut microbiome composition.

## 6. Methods

### 6.1. Study design and population

This prospective, randomized, double-blind, placebo-controlled trial was conducted with 26 West-Highland terriers, 3 Scottish Terriers, and one Cairn Terrier, bred in a single facility in France. The study was approved by the institutional ethics committee (Saisine 2024-06-29, Ecole Nationale Vétérinaire d’Alfort, Maisons-Alfort, France) and conducted in accordance with national and institutional regulations for animal experimentation.

All dogs enrolled in the study were confirmed to be naturally infected with *G. duodenalis*. Inclusion criteria were generally good health status; presence of acute or chronic diarrhea (watery or soft feces); and coproscopic confirmation of *Giardia* infection. Exclusion criteria were identification of a significant systemic disease other than giardiasis; refusal of the proposed dietary regimen; administration of any treatment within 15 days prior to inclusion; and administration of any pre-or postbiotic treatment within one month prior to inclusion.

Animals were randomly allocated at day −1 (D−1) to two equally sized groups (n = 15 per group) (**Figure 1**): a placebo group receiving metronidazole plus a tablet with placebo or an LBHF group receiving metronidazole plus a tablet containing the LBHF postbiotic (one tablet per day). Randomization was performed using a sealed-envelope online randomization tool (https://www.sealedenvelope.com/). Groups were balanced on body weight and age.

**Figure 1.**
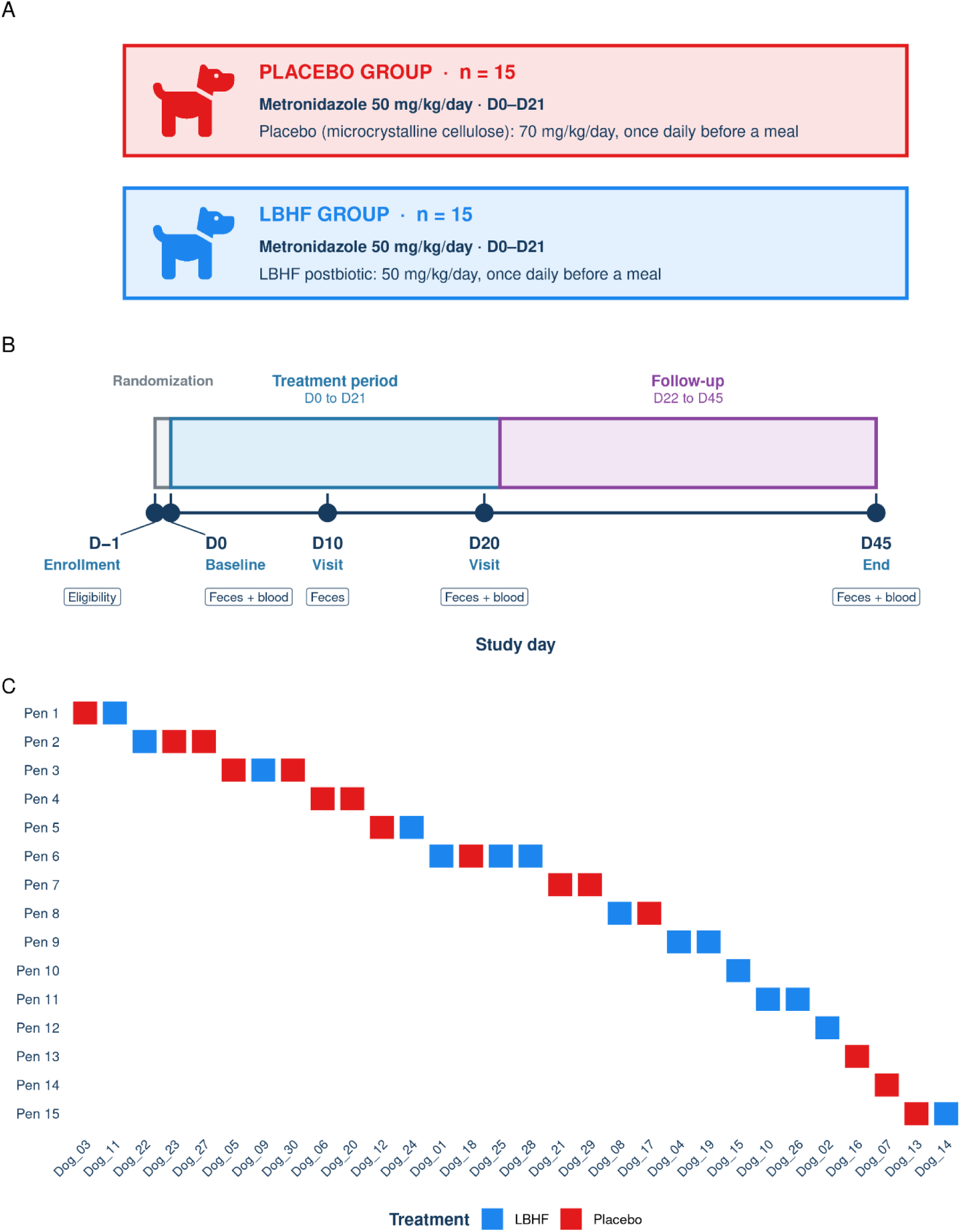
Study protocol overview. (A) Treatment group composition. (B) Study timeline, phases, and collection timepoints. (C) Pen structure of the study.

### 6.2. Dietary standardization and treatment protocol

The study comprised two sequential phases (**Figure 1**): a 3-week treatment period (D0–D21) followed by a 3-week follow-up period (D22– D45). Throughout the study, animals were fed a commercially available extruded diet (PRO PLAN® Medium Puppy Sensitive Digestion, Lamb, Purina, France). The diet did not contain any added probiotics, prebiotics, or postbiotics.

From D0 to D21, all animals received the standard antiparasitic treatment for canine giardiasis, metronidazole (Metrobactin®, Dechra, France), at 50 mg/kg/day administered orally once daily. In addition, animals either received a tablet with placebo (microcrystalline cellulose; MCC) or with LBHF. The tablets were prepared by Synergie Prod SAS (Sevremoine, France) and further mainly contained dicalcium phosphate, with some pig liver extract as palatant and colorant, silicic acid, and magnesium stearate. Tablets were visually indistinguishable and were packaged in bottles with codes so that the content was only known to study sponsor. The intent was to treat each dog 600 mg LBHF per day.

### 6.3. Clinical monitoring and sample collection schedule

Clinical examinations were performed at four timepoints: D0, D10, D20, and D45. Stool samples (minimum 10 g) were collected at D-1, D0, D10, D20 and D45 for parasitological analyses and for microbiological analyses at D0 and D45. A total of 4 mL of whole blood was collected in a serum separator tube (BD Vacutainer® SST^TM^ II ADVANCE, BD-Belliver Industrial Estate, Belliver Way, Roborough, Plymouth UK) to obtain serum at D0 and D45 for biochemical and immunological analyses.

### 6.4. *Giardia* cyst quantification

At inclusion (D-1), and throughout trial period (D0, D10, D20, and D45), *Giardia* cyst counts were quantified using direct immunofluorescence assays (IFA) (MeriFluor Cryptosporidium/Giardia, Meridian Bioscience Europe, Milan, Italy) as per the manufacturer’s instructions with modifications as reported previously (Mammeri et al., 2019). Results are expressed as cysts per gram of feces.

### 6.5. Fecal variables

At each visit, fecal consistency was scored using the 5-point Bristol stool scale (BSS) adapted for dogs: 1 = watery (very liquid, flows from container); 2 = soft, unformed (takes shape of container); 3 = soft, formed, moist (retains shape); 4 = firm, formed, dry (remains firm and moist); 5 = hard dry pellets (Carciofi et al., 2009; Félix et al., 2013; Murakami et al., 2018). Fecal odor was scored on a 3-point scale (1 = no odor, 2 = moderate, 3 = strong) but not on day -1. The scoring was based on previously reported 2, 5 or 10-point Likert scales (Beynen C Saris, 2014; Dos Reis et al., 2016; Jeusette et al., 2025) used in studies on fecal odor in dogs and a 3 point scale was deemed most useful in the facility in which our study was performed. Fecal color was noted when abnormal (e.g., presence of fresh blood, yellow diarrhea). One-gram aliquots of feces were stored at −80°C for subsequent quantification of fecal calprotectin and fecal secretory IgA (sIgA), both performed by an external laboratory (Laboklin, France).

The canine dysbiosis index (CDI) was determined at each timepoint using quantitative PCR targeting seven bacterial taxa: *Faecalibacterium*, *Turicibacter*, *Streptococcus*, *Escherichia coli*, *Blautia*, *Fusobacterium*, and *Peptacetobacter hiranonis,* (formerly *Clostridium*); as previously described (AlShawaqfeh et al., 2017), which was performed by IDEXX Laboratories (France).

### 6.6. Blood biomarkers

Blood serum samples were collected at D0 and D45 for quantification of markers of digestive function, including serum IgA (mg/mL; IgA ELISA Kit ABIN7041551, detection range 31.25–1000 ng/mL), vitamin B12 (cobalamin, pg/mL, Immulite 2000 Xpi, Siemens), vitamin B9 (folate, ng/mL, Immulite 2000 Xpi, Siemens), canine trypsin-like immunoreactivity (TLI, ng/mL, Immulite 2000 Xpi, Siemens), pancreatic lipase activity (DGGR method, U/L, Cubevet, Eurolyser), and canine cortisol (nmol/L, Immulite 2000 Xpi, Siemens). All analyses were performed at the BioPôle analytical laboratory (EnvA, Maisons-Alfort, France). The feces-to-blood IgA ratio was computed from matched fecal sIgA and serum IgA measurements.

### 6.7. Metagenomic shotgun sequencing and bioinformatic processing

#### 6.7.1. DNA Extraction Methods

At baseline (D0) and day 45 (D45), 1-g fecal aliquots were collected and transferred within one hour of collection into OMNIgene®•GUT collection tubes (DNA Genotek™, OraSure Technologies Inc., Ottawa, Canada). Samples were subsequently stored at −80°C until batch shipment.DNA from samples was isolated using the QIAGEN DNeasy PowerSoil Pro Kit, according to the manufacturer’s protocol. Isolated DNA was quantified using Qubit Flex fluorometer and Qubit™ dsDNA HS Assay Kit (Thermofisher Scientific).

#### 6.7.2. Library Preparation and Sequencing Methods

DNA libraries were prepared using the Watchmaker DNA Library Prep Kit (7K0019-1K). Genomic DNA was fragmented using a mastermix of Watchmaker Frag/AT Buffer and Frag/AT Enzyme Mix. IDT xGen UDI Primers and IDT Stubby Adapters were added to each sample followed by 7 cycles of PCR to construct the DNA libraries. The final DNA libraries were purified using CleanNGS magnetic beads (CleanNA) and eluted in nuclease-free water. Following elution, the libraries were quantified using the Qubit™ fluorometer dsDNA HS Assay Kit. Libraries were circularized and sequenced on the Element AVITI platform using the 2×150 Cloudbreak FreeStyle sequencing kit.

#### 6.7.3. Taxonomic and functional quantification

Sequencing files were processed as previously described (Eckhardt et al., 2026). Reads underwent quality control (fastp v0.20.1 (Chen, 2025): length > 15 bp, ambiguous reads < 5, percentage of reads below Q15 < 40%, sequence complexity > 30%), prior to host-read decontamination by mapping to the Canis familiaris reference genome with RefSeq accession ‘GCF_011100685.1’. Kraken2 (v2.07) (Wood et al., 2019) on a 0.1 confidence threshold and Bracken (v3.0) (Lu et al., 2017) were used for taxonomic profiling on a previously described internal genomic database (Eckhardt et al., 2026). A minimum of 10 Bracken counts were required to consider a taxon as present.

Sequencing reads were further mapped against a previously published dog gut metagenomic gene catalog of 1,247,405 genes, predicted on genomes also included in the taxonomic database (Coelho et al., 2018). Reads were mapped into the catalog using BWA mem (v0.7.17) (Li, 2013), and counts were extracted with SAMtools (v1.20) (Danecek et al., 2021). Gene counts were normalized as transcripts per million (TPM). Finally, we grouped genes with the same CAZyme annotation (dbCAN2 v1.1) (Zhang et al., 2018) to produce the final CAZyme abundance table, and into KEGG Orthology (KO) groups, annotated in the original publication (Coelho et al., 2018).

### 6.8. Statistical analyses

#### 6.8.1. Endpoint statistical testing

Left- and right-censored assay values were handled by single-value substitution, using 50% of the reported detection threshold for lower-censored observations and 110% of the threshold for upper-censored observations. Baseline group differences (D0) were assessed using Wilcoxon rank-sum tests for continuous endpoints and Fisher’s exact test for sex composition. P-values were adjusted for multiple comparisons across all baseline endpoints using the Benjamini–Hochberg false discovery rate (BH FDR) procedure.

#### 6.8.2. Temporal analysis of the primary endpoint

*Giardia* cyst counts were inverse hyperbolic sine (asinh) transformed prior to modelling. The individual-level baseline covariate was defined as the mean of the day −1 and day 0 measurements per animal. A linear mixed-effects ANCOVA model was fitted to post-treatment observations (days 10, 20, and 45) using the nlme package in R. The fixed-effects structure was:

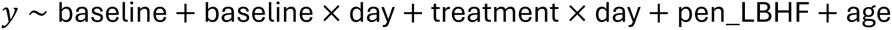

where *day* was treated as a categorical factor, *baseline* denotes the animal-level pre-treatment mean, and *pen_LBHF* is the number of LBHF-assigned animals in the same pen (excluding the focal animal), included to account for potential pen-level contamination effects. Random intercepts were included for both the individual animal and the pen. Models were fitted by REML. To obtain a single overall treatment effect estimate, a reduced model without the treatment × day interaction was additionally fitted; the treatment coefficient from this model is reported alongside its standard error and P-value. Per-day marginal means and pairwise treatment contrasts were obtained using the emmeans v2.03 R package (Lenth C Piaskowski, n.d.).

#### 6.8.3. Temporal analyses of secondary endpoints

**Fecal endpoints** (BSS, fecal odor score, fecal calprotectin, fecal sIgA, and canine dysbiosis index) were measured at days 0, 10, 20, and 45 (BSS scoring was also available at day −1). Each endpoint was modelled with a longitudinal ANCOVA using the same random-effects structure as the primary endpoint (random intercepts for animal and pen). The fixed-effects structure was:

The individual-level baseline was defined as the mean of available day −1 and day 0 values. Unlike the primary endpoint model, the treatment × day interaction was not included for secondary endpoints; a single treatment coefficient is reported for each. Models were restricted to post-treatment timepoints (days ≥ 10).

**Blood biomarkers** (pancreatic lipase, vitamins B12 and B9, trypsin-like immunoreactivity, cortisol, IgA, and feces-to-blood IgA ratio) were measured at days 0 and 45 only. These were analyzed with a cross-sectional ANCOVA comparing the day-45 value to baseline (mean of days −1 and 0), with pen as a random effect and age and pen_LBHF as additional fixed covariates:

All blood biomarker concentrations were log_10_-transformed prior to modeling. P-values from all secondary endpoint models were adjusted jointly using the BH FDR procedure.

#### 6.8.4. Distance-based and community-level analysis

Aitchison distances between species-level gut microbiome compositional profiles were computed after centered log-ratio (CLR) transformation, using half the minimum non-zero value per species as a pseudocount. Multivariable PERMANOVA was fitted using adonis2 (vegan v2.7.3) (Oksanen, 2007) with all covariates entered simultaneously, reporting marginal effects; univariable models were fitted separately for each covariate. P-values were computed with 999 permutations. The interaction of treatment group and study day was used as the primary test of treatment effect on community composition.

To assess within-subject temporal stability, we computed pairwise Aitchison distances between day 0 and day 45 samples for each animal. The observed difference in mean within-subject distance between treatment groups was tested against a null distribution generated from 1,000 within-pen label permutations (i.e., treatment labels were shuffled only among animals sharing a pen). A complementary analysis compared the mean within-subject distance to the mean between-subject distance across all unrelated pairs, using a permutation test without pen restriction. These analyses were repeated on CLR-transformed CAZyme abundances.

#### 6.8.5. Multivariate assessment of treatment effect on microbial communities

CLR-transformed species, KO, and CAZyme abundances were baseline-normalised by subtracting each animal’s day-0 value from its day-45 value prior to PLS-DA, so that the model discriminates on treatment-associated compositional change rather than baseline state. PLS-DA was fitted using mixOmics (v6.34.0) (Rohart et al., 2017) with two components, no feature scaling, and either all prevalent features (non-sparse model) or a sparse model retaining 10 features per component (keepX = 10). Classification performance was estimated by leave-one-out cross-validation, reporting the balanced error rate (BER) under centroid-distance classification. Component loadings were extracted to rank individual feature contributions.

Gene-set enrichment analysis of PLS-DA loading vectors was performed using fgsea (v1.36.2) (Korotkevich et al., 2021). Species loadings were enriched against a hierarchical taxonomy table spanning genus through phylum; KO loadings were enriched against KEGG pathway annotations; CAZyme loadings were enriched against substrate categories from the CAZy database hierarchy. Enrichment was run with 1,000 permutations, a minimum gene-set size of 5, and a maximum of 500. Normalized enrichment scores (NES) and BH FDR are reported.

#### 6.8.6. Microbiome univariate association analysis

CLR-transformed microbial feature abundances (species, KO terms, and CAZyme families) at the follow-up timepoint (day 45) were modelled independently as a function of treatment group using linear mixed-effects ANCOVA, implemented with nlme v3.1.168 (REML). The fixed-effects structure was:

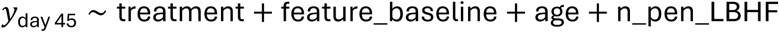

where *feature_baseline* is the CLR-transformed abundance of the same feature at day 0, *age* is the animal’s age in days, and *n_pen_LBHF* is the number of LBHF-assigned animals in the same pen (excluding the focal animal). A random intercept for pen was included to account for shared pen environment. Features were retained for modelling if they met layer-specific prevalence and abundance thresholds: species required prevalence ≥ 20% and mean relative abundance ≥ 1 × 10^−4^; KO terms required prevalence ≥ 30% and mean relative abundance ≥ 1 × 10^−4^; CAZyme families required prevalence ≥ 20% with no minimum abundance threshold. P-values for the treatment term were adjusted using the H FDR procedure within each data layer.

Species-level Shannon diversity was computed on raw relative abundances using the vegan package and modelled with the same ANCOVA framework, substituting the species-level diversity at day 0 as the baseline covariate:

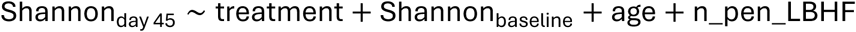

A random intercept for pen was retained.

#### 6.8.7. Association of microbiome species with *Giardia* burden

Species-level CLR-transformed abundances were tested for association with asinh-transformed *Giardia* cyst counts using a linear model for baseline samples (day 0). The structure was:

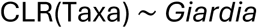

where *Giardia* is the asinh-transformed cyst count. Species were retained for modelling if they met a prevalence cut-off of ≥ 30%. P-values for the *Giardia* term were adjusted using the BH FDR procedure across all tested species. Taxonomic enrichment of the full t-statistic ranking was performed using fgsea against the species taxonomy hierarchy (family and phylum levels), with a minimum set size of 3.

The same model was used to associate Shannon diversity baseline levels with *Giardia* cyst counts.

#### 6.8.8. *Giardia* counts prediction using taxonomic abundance

CLR-transformed baseline (day 0) species abundances were used to predict asinh-transformed *Giardia* cyst counts. We performed leave-one-out cross-validation to assess the prediction performance. A default (nrounds = 100, objective = reg:squarederror) XGBoost model was fitted on the complete dataset minus one sample at a time (R xgboost v3.2.1.1 package). Predicted *Giardia* cyst counts in left-out samples were correlated with the observed concentration values by means of a Spearman correlation.

Feature importance was assessed by means of SHapley Additive exPlanations (SHAP) values (shapviz v0.10.3) (Lundberg C Lee, 2017). Per feature, the average absolute SHAP value was computed as an importance statistic. Directionality was assessed correlating the input abundance values to their respective SHAP values by means of a Spearman correlation.

## 7. Results

### 7.1. Study population and giardiasis

A total of 30 juvenile dogs were enrolled in this study (median age 2 years, interquartile range (IQR) = 3.75 years). All animals had confirmed natural *Giardia* infection at the time of enrollment (*Giardia* cysts/g feces: median = 8,000, IQR = 29,000, max = 308,000). Animals were randomly allocated to two equally-sized groups (placebo; n = 15 and LBHF; n = 15) with comparable *Giardia* cyst load (Wilcoxon rank-sum test, mean_placebo_ = 48,767 opg, mean_LBHF_ = 24,633 opg, P = 0.47), comparable ages (mean_placebo_ = 3.35 years, mean_LBHF_ = 3.05 years, P = 0.77), and comparable sex composition (73% vs. 53% female, Fisher’s exact test P = 0.45) at the time of randomization (day −1). Both groups received conventional treatment for *Giardia* infection (metronidazole) starting at day 0. In addition to metronidazole, the LBHF group received on average 50 mg / kg /day of LBHF supplementation with a single tablet per day.

As dogs could not be weighed in the facility, but the age of each dog was recorded (Table 1), the dose was obtained by using bodyweights estimated based on growth curves of West Highland Terriers. The daily dose of 50 mg / kg / days was a downwards revision from the intended dose in the protocol because flow cytometry analysis of the tablets revealed that only 420 mg LBHF was present (2.5*10^10^ cell bodies instead of the intended 3.6*10^10^).

**Table 1.** Between-group comparison at baseline (day 0), before intervention or metronidazole administration. Continuous variables: means ± SD, compared by Wilcoxon rank-sum test. Sex composition: proportion female, compared by Fisher’s exact test. FDR correction applied across all endpoints.

| Category | Endpoint | N | Placebo | LBHF | P | FDR |
|---|---|---|---|---|---|---|
| Anthropometric | Age (years) | 30 | 3.35 ± 3.49 | 3.05 ± 2.27 | 0.770 | 0.976 |
| Anthropometric | Sex (prop. female) | 30 | 0.73 | 0.53 | 0.450 | 0.750 |
| Primary | Giardia cysts/g feces | 30 | 45,700 ± 40,637 | 7,067 ± 6,619 | 2.83e-04 | 0.002 |
| Fecal | Fecal score (BSS) | 30 | 2.97 ± 0.74 | 3.53 ± 0.44 | 0.028 | 0.098 |
| Fecal | Fecal odor score | 30 | 2.47 ± 0.52 | 1.47 ± 0.52 | 1.49e-04 | 0.002 |
| Fecal | Fecal calprotectin (µg/g) | 30 | 73.73 ± 53.62 | 116.20 ± 112.58 | 0.372 | 0.652 |
| Fecal | Fecal sIgA (mg/g) | 30 | 3.52 ± 1.95 | 4.39 ± 2.17 | 0.313 | 0.652 |
| Blood | Pancreatic lipase DGGR (U/L) | 28 | 59.58 ± 30.54 | 55.26 ± 19.49 | 1.000 | 1.000 |
| Blood | B12 (pg/mL) | 28 | 727.79 ± 343.08 | 765.86 ± 265.23 | 0.836 | 0.976 |
| Blood | B9 (ng/mL) | 27 | 19.66 ± 5.29 | 19.63 ± 4.65 | 0.942 | 1.000 |
| Blood | Trypsin-like immunoreactivity (ng/mL) | 27 | 27.29 ± 10.54 | 22.53 ± 7.68 | 0.216 | 0.605 |
| Blood | Cortisol (nmol/L) | 28 | 29.51 ± 22.95 | 25.03 ± 16.75 | 0.760 | 0.976 |
| Blood | IgA (mg/mL) | 29 | 4.49 ± 1.61 | 5.03 ± 1.82 | 0.371 | 0.652 |
| Ratio | Feces-to-blood IgA ratio | 20 | 2.04 ± 0.99 | 2.45 ± 1.43 | 0.761 | 0.976 |
| Bacterial | Canine Dysbiosis Index | 30 | −0.63 ± 2.27 | −2.94 ± 2.03 | 0.003 | 0.016 |

At the time of treatment initiation (day 0), *Giardia* cyst load had diverged between groups (mean_placebo_ = 45,700, mean_LBHF_ = 7,067, P = 2.83 × 10^−4^), suggesting differential pre-treatment infection burden. Additional baseline differences at day 0 included fecal odor score (mean_placebo_ = 2.47, mean_LBHF_ = 1.47, P = 1.49 × 10^−4^) and canine dysbiosis index (mean_placebo_ = −0.633, mean_LBHF_ = −2.94, P = 3.43 × 10^−3^), with BSS showing a non-significant trend toward higher scores in the LBHF group (mean_placebo_ = 2.97, mean_LBHF_ = 3.53, P = 0.098). No significant between-group differences were observed in blood biomarkers or sex composition at baseline (**Table 1**).

### 7.2. Association between LBHF treatment, *Giardia* clearance, and fecal parameters

Given the baseline imbalance in *Giardia* burden between groups (see above), we used a longitudinal ANCOVA model adjusting for individual baseline levels and their interaction with time, with pen and age as additional covariates (see Methods). *Giardia* cyst counts were inverse hyperbolic sine (asinh) transformed prior to analysis.

The LBHF group showed markedly lower *Giardia* concentrations across the entire treatment period (effect of placebo relative to LBHF: estimate = 6.92, SE = 0.71, P = 8.71 × 10^−10^; **Figure 2A**). This difference was already evident at day 10, the first post-treatment timepoint (marginal mean_LBHF_ = 0.11, marginal mean_placebo_ = 10.3, P = 2.16 × 10^−10^), and remained large at day 20 (marginal mean_LBHF_ = 1.98, marginal mean_placebo_ = 10.2, P = 1.49 × 10^−8^). By day 45, *Giardia* concentrations in the placebo group had also decreased substantially, and the between-group difference was no longer statistically significant (marginal mean_LBHF_ = 0.58, marginal mean_placebo_ = 2.26, P = 0.12).

**Figure 2.**
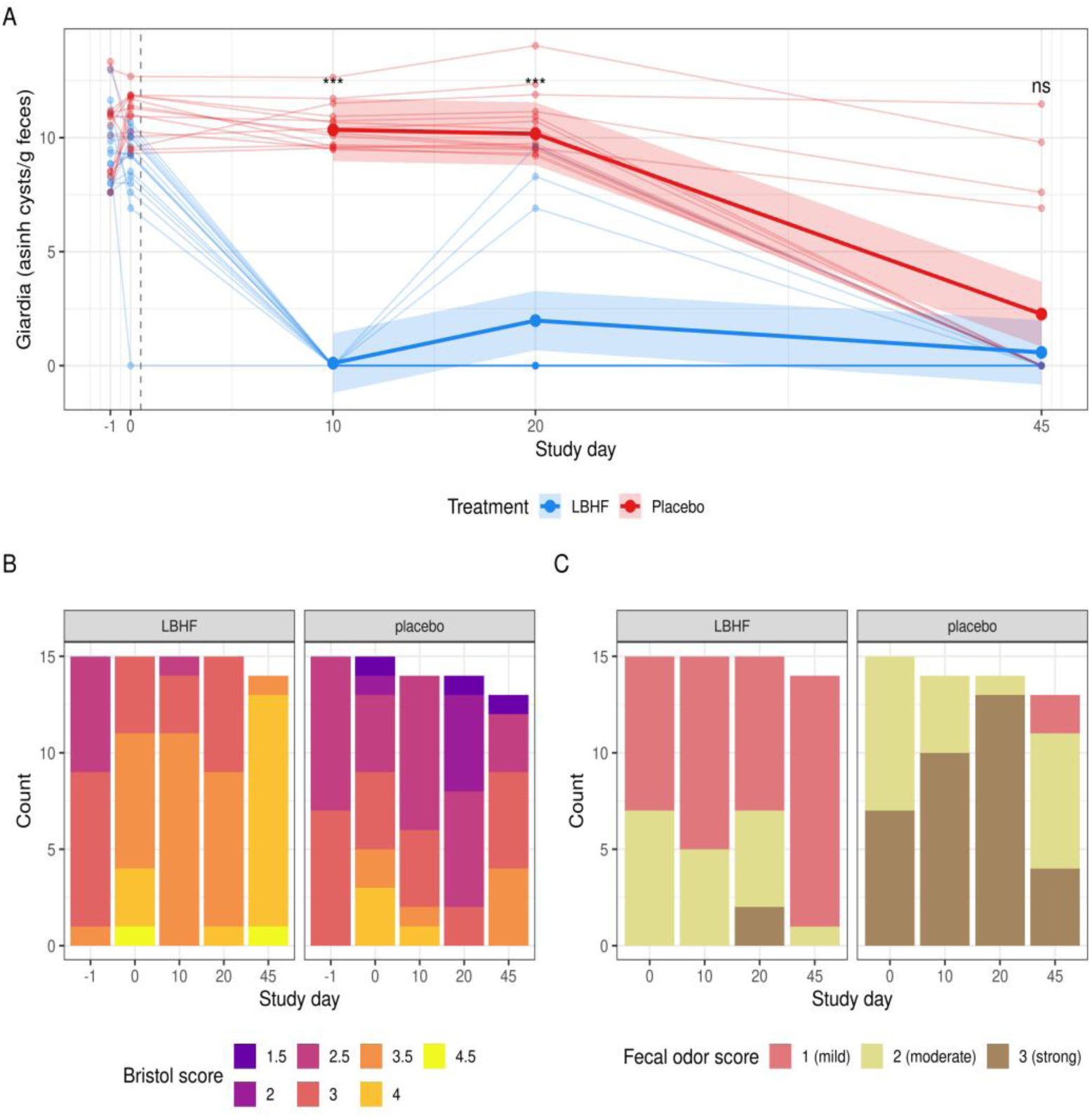
LBHF treatment accelerates Giardia clearance and improves fecal parameters. (A) Individual Giardia trajectories (thin lines, asinh-transformed cysts/g feces) across all timepoints, with model-estimated marginal means ± S5% CI (thick lines and shaded ribbons) from the longitudinal ANCOVA overlaid for post-treatment days (10, 20, 45). The dashed vertical line marks the start of treatment (day 0). Significance annotations reflect per-day pairwise contrasts between groups (*** P < 0.001; ns, not significant). Blue: LBHF; Red: Placebo. (B) Distribution of BSS scores by study day and treatment group. (C) Distribution of fecal odor scores by study day and treatment group.

We next assessed whether LBHF also influenced fecal parameters (BSS, fecal odor score, fecal calprotectin, fecal sIgA, and canine dysbiosis index), measured at four timepoints (days 0, 10, 20, and 45; BSS also included day −1), and blood biomarkers (measured at days 0 and 45). Longitudinal ANCOVA models were used for fecal endpoints; ANCOVA comparing baseline to end-of-study was used for blood endpoints. Two fecal outcomes improved significantly with LBHF treatment: BSS increased toward healthier stool consistency (estimate = 0.75, P = 1.08 × 10^−7^, FDR = 6.46 × 10^−7^; **Figure 2B**) and fecal odor score decreased (estimate = −1.28, P = 2.38 × 10^−8^, FDR = 2.86 × 10^−7^; **Figure 2C**). In the LBHF group, the modal BSS progressed from 3.5 at day 0 to 4.0 at day 45; in the placebo group it moved from 2.5 to 3.0. No significant treatment effects were detected for dysbiosis index (estimate = −1.42, P = 0.16), fecal calprotectin, fecal sIgA, or any blood biomarker after FDR correction (Table 2), although dysbiosis, calprotectin and sIgA showed a trend for lower values in the LBHF treated group.

**Table 2.** Treatment effect (LBHF vs. placebo) on fecal and blood biomarkers. Estimates reflect the LBHF effect (positive = increase in LBHF group). Longitudinal ANCOVA models were used for fecal endpoints (days 0, 10, 20, 45); ANCOVA comparing day 0 to day 45 was used for blood endpoints. FDR correction applied across all endpoints.

| Endpoint | N | Estimate | P-value | FDR |
|---|---|---|---|---|
| Fecal odor score | 85 | −1.276 | 2.38e-08 | 2.86e-07 |
| Fecal score (BSS) | 85 | 0.749 | 1.08e-07 | 6.46e-07 |
| Fecal calprotectin (µg/g) | 85 | −0.331 | 1.08e-01 | 3.89e-01 |
| Canine Dysbiosis Index | 85 | −1.420 | 1.57e-01 | 3.89e-01 |
| Fecal sIgA (mg/g) | 85 | −0.109 | 1.73e-01 | 3.89e-01 |
| Blood IgA (mg/mL) | 27 | 0.067 | 1.94e-01 | 3.89e-01 |
| Pancreatic lipase DGGR (U/L) | 26 | 11.304 | 2.81e-01 | 4.82e-01 |
| Blood B12 (pg/mL) | 26 | −0.031 | 4.27e-01 | 6.40e-01 |
| Feces-to-blood IgA ratio | 13 | 0.128 | 5.20e-01 | 6.93e-01 |
| Trypsin-like immunoreactivity (ng/mL) | 24 | −0.042 | 5.86e-01 | 7.03e-01 |
| Blood B9 (ng/mL) | 25 | −0.023 | 6.69e-01 | 7.30e-01 |
| Blood cortisol (nmol/L) | 26 | 0.023 | 8.55e-01 | 8.55e-01 |

### 7.3. Microbial community-level rearrangements in response to LBHF treatment

Metagenomic shotgun sequencing was performed for all puppies at baseline (day 0) and for 27 of them at the last study timepoint (day 45). The species-level compositional profiles were obtained (see Methods) and transformed using centered log-ratio (CLR) prior to analysis. To assess overall temporal community change, we calculated within-subject Aitchison distances between the day 0 and day 45 samples. Both treatment groups showed substantial microbiome turnover over the intervention period, and the magnitude of change did not differ significantly between groups (mean Aitchison distance, LBHF = 32.8 ± 7.37, placebo = 31.3 ± 8.7; Wilcoxon rank-sum test, P = 0.52) (Figure 3**A**). As expected, within-subject samples were considerably more similar to each other than to samples from different animals (mean distance, same animal = 32.1 vs. different animal = 38.5; permutation test, P < 0.001), confirming a strong individual signature in microbiome composition.

**Figure 3.**
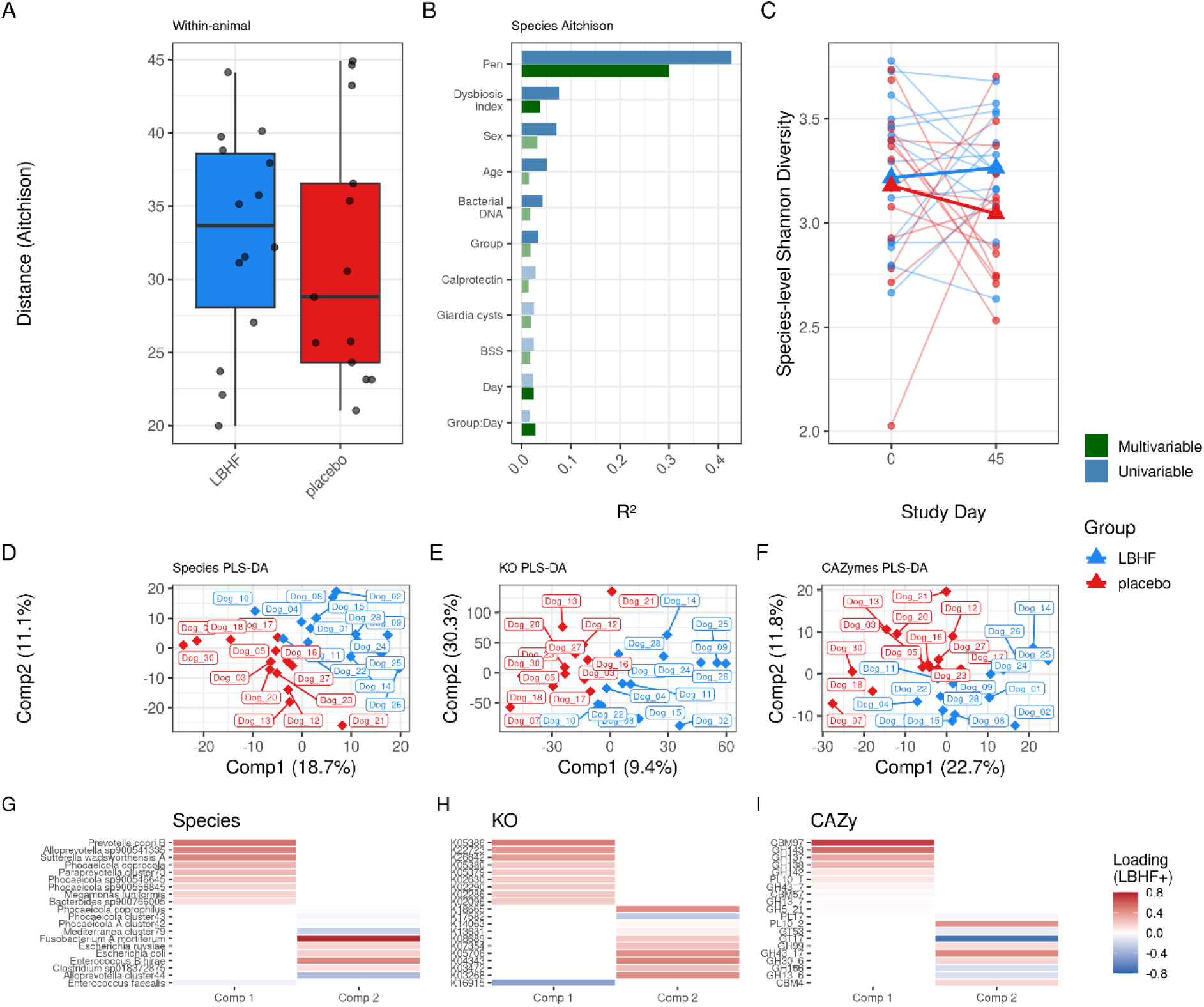
(A) Within-subject Aitchison distances between day 0 and day 45 samples, computed from CLR-transformed Kraken/Bracken species-level abundances, shown by treatment group. (B) Marginal R² values from multivariable PERMANOVA on the species-level Aitchison distance matrix for each explanatory variable (treatment group, study day, their interaction, and clinical covariates: canine dysbiosis index, total bacterial DNA, fecal calprotectin, fecal score, Giardia cysts, sex, and age). (C) Species-level Shannon diversity over time by treatment group; thin lines represent individual animals and thick lines the group mean. (D–F) PLS-DA score plots for baseline-normalized CLR-transformed species (D), KEGG Orthology (KO) functional (E), and CAZyme family (F) profiles; percentages on axes indicate variance explained by each component. LBHF animals are shown in blue, placebo in red. (G–I) Sparse PLS-DA loading heatmaps (top 10 features per component) for species (G), KO terms (H), and CAZy families (I). Loading values are oriented so that positive values (red) correspond to enrichment in LBHF-treated animals relative to placebo.

To identify which host and clinical covariates explained interindividual variation in species-level composition, we fitted multivariable PERMANOVA models on the Aitchison distance matrix (marginal effects) Figure 3**B**). The dominant source of variation was pen membership (R^2^ = 0.31, P < 0.001), reflecting not only shared environment effect, but also including the individuality of each animal in time. Beyond pen, significant marginal contributions were found for the canine dysbiosis index (R^2^ = 0.040, P = 0.003), total bacterial DNA abundance (R^2^ = 0.023, P = 0.025), study day (R^2^ = 0.025, P = 0.019), and treatment group (R^2^ = 0.022, P = 0.045). Notably, the interaction of treatment group with study day was also significant (R^2^ = 0.024, P = 0.028), indicating that the microbiome trajectories of the two treatment groups diverged over the course of the study beyond any difference in starting composition.

We then assessed whether LBHF treatment was associated with changes in alpha diversity, measured by species-level Shannon diversity at days 0 and 45 (Figure 3**C**). Overall, Shannon diversity was similar between groups at baseline (mean_LBHF_ = 3.22 ± 0.37, mean_placebo_ = 3.18 ± 0.44) and showed a modest, non-significant trend toward higher diversity in the LBHF group over the course of the study (ANCOVA, estimate_LBHF_ = 0.22, P = 0.13; mean_LBHF_ at day 45 = 3.26 ± 0.28, mean_placebo_ at day 45 = 3.04 ± 0.34).

### 7.4. LBHF treatment remodels the gut microbiome toward increased Bacteroidota representation and enhanced capacity for plant polysaccharide degradation

To characterize which microbial species, functional genes (KEGG orthologues, KO), and carbohydrate-active enzymes (CAZymes) were differentially altered by LBHF treatment, we applied complementary multivariate and univariate analytical frameworks. The multivariate approach used Partial Least Squares Discriminant Analysis (PLS-DA) to identify axes of joint community variation discriminating between treatment groups, whereas the univariate approach tested each feature independently using linear mixed-effects ANCOVA.

For the multivariate analysis, each data layer was baseline-normalized prior to modelling by subtracting each animal’s day-0 value from its day-45 value, so that the discriminating axes capture treatment-associated *change* in composition rather than pre-existing differences. For each layer, we fitted two complementary models: a non-sparse model, in which all prevalent features are retained, enabling gene-set enrichment analysis of the full loading vectors; and a sparse model (10 features per component), which concentrates the discriminating axes on the smallest feature subset that maximizes group separation. Discriminatory performance was estimated by leave-one-out cross-validation (LOOCV), with the balanced error rate (BER) under centroid-distance classification used as the primary metric (BER = 0.5 corresponds to chance-level discrimination).

At the species level, the non-sparse model achieved only modest discrimination in component 1 (BER = 0.48); restricting the sparse model to 10 features per component substantially improved performance (BER = 0.33) (Figure 3**D**). Enrichment analysis of the full component 1 loading vector identified a consistent taxonomic signal: LBHF-associated community change was characterized by relative increases in Bacteroidales (NES = 2.61, FDR = 2.71 × 10^−10^), Burkholderiales (NES = 1.98, FDR = 9.28 × 10^−4^), *Phocaeicola* (NES = 1.90, FDR = 4.03 × 10^−3^), and *Sutterella* (NES = 1.82, FDR = 2.09 × 10^−3^), and relative decreases in Bacilli (NES = −2.01, FDR = 1.08 × 10^−4^), Lactobacillaceae (NES = −1.93, FDR = 5.05 × 10^−3^), Campylobacterales (NES = −1.76, FDR = 4.02 × 10^−2^), and Clostridia (NES = −1.70, FDR = 5.05 × 10^−3^) Accordingly, the species with the highest sparse loadings on component 1 were *Prevotella copri* B, *Alloprevotella* sp., and *Sutterella wadsworthensis* A, the first two belonging to Bacteroidota/Bacteroidales and the latter to Pseudomonadota/Burkholderiales, consistent with the enrichment results (Figure 3**G**).

Beyond taxonomy, we examined the functional gene repertoire encoded by the microbiome by profiling KEGG Orthologues (KOs): a catalogue of protein-coding genes grouped by function, spanning metabolic, signaling, and structural pathways. Applied to the same baseline-normalized framework, the KO-level PLS-DA revealed that the treatment-associated functional signal was distributed across two partially orthogonal axes (non-sparse BER comp1 = 0.52, comp2 = 0.34; sparse BER comp1 = 0.45, comp2 = 0.67) (Figure 3**E**). Enrichment of component 1 loadings (Figure 3**H**) pointed to depletion in LBHF treated animals of teichoic acid biosynthesis KOs (NES = −1.88, FDR = 0.021), which are predominantly found in Gram-positive bacteria, and *Helicobacter pylori* epithelial-cell-signaling KOs (NES = −1.98, FDR = 0.021) (**Table 3**). Component 2 showed further depletion of KO sets associated with motility and colonization — flagellar assembly (NES = −2.31, FDR < 0.001), biofilm formation (NES = −1.84, FDR < 0.001), quorum sensing (NES = −1.46, FDR = 0.029), and two-component signaling (NES = −1.32, FDR = 0.042); in addition to an enrichment for lipoarabinomannan biosynthesis KOs (NES = 2.16, FDR = 0.039) (**Table 3**), potentially reflecting increased representation of Actinobacteria-associated cell-envelope features.

**Table 3.** Set enrichment analysis results (FDR < 0.05) of fecal microbiome gene abundances (KO and CAZymes) impacted by LBHF (vs. placebo) treatment. Feature loads on two-component PLS models were used for ranking. KO categories correspond to KEGG pathways, while CAzyme ontology corresponds to substrate classes.

| Feature | pval | padj | NES | size | component | Class |
|---|---|---|---|---|---|---|
| map02040: Flagellar assembly | 8.13e-11 | 2.07e-08 | -2.33 | 54 | comp2 | KO |
| map03010: Ribosome | 5.18e-07 | 4.38e-05 | 2.48 | 58 | comp2 | KO |
| map05111: Biofilm formation | 5.38e-06 | 3.42e-04 | -1.85 | 74 | comp2 | KO |
| map01052: Type I polyketide structures | 1.28e-04 | 6.50e-03 | -2.00 | 10 | comp2 | KO |
| map02024: Quorum sensing | 6.00e-04 | 2.54e-02 | -1.45 | 188 | comp2 | KO |
| map00571: Lipoarabinomannan (LAM) biosynthesis | 7.32e-04 | 2.57e-02 | 2.07 | 12 | comp2 | KO |
| map02020: Two-component system | 9.11e-04 | 2.57e-02 | -1.31 | 440 | comp2 | KO |
| map04624: Toll and Imd signaling pathway | 9.09e-04 | 2.57e-02 | -1.91 | 6 | comp2 | KO |
| map05120: Epithelial cell signaling in Helicobacter pylori infection | 2.51e-04 | 3.19e-02 | -1.98 | 13 | comp1 | KO |
| Pectin | 1.88e-07 | 5.09e-06 | 1.82 | 98 | comp1 | CAZy |
| Plant | 2.53e-05 | 3.42e-04 | 1.56 | 228 | comp1 | CAZy |
| NSP | 6.04e-05 | 5.44e-04 | 1.52 | 209 | comp1 | CAZy |
| DF | 3.03e-04 | 2.05e-03 | 1.47 | 260 | comp1 | CAZy |
| Hemicellulose | 1.21e-03 | 5.44e-03 | 1.55 | 89 | comp1 | CAZy |
| Structural | 1.10e-03 | 5.44e-03 | 1.44 | 267 | comp1 | CAZy |
| GAG | 3.26e-04 | 8.79e-03 | 1.98 | 36 | comp2 | CAZy |
| Fructan | 4.87e-03 | 1.88e-02 | -1.82 | 6 | comp1 | CAZy |

We additionally profiled carbohydrate-active enzymes (CAZymes): a specialized gene catalogue encoding enzymes that synthesize, degrade, and modify complex carbohydrates, which provide a direct readout of the microbiome’s capacity to process dietary and host-derived polysaccharides. The CAZyme layer yielded the most discriminatory models overall (non-sparse BER comp1 = 0.44, comp2 = 0.41; sparse BER comp1 = 0.30, comp2 = 0.33) (Figure 3**F**). Enrichment analysis indicated that LBHF-associated community change was accompanied by increased capacity for degradation of plant-derived polysaccharides (**Table 3**), including pectin (NES = 1.86, FDR = 6.3 × 10^−6^), hemicellulose (NES = 1.57, FDR = 6.6 × 10^−3^), and non-starch polysaccharides (NES = 1.54, FDR = 1.1 × 10^−3^), and decreased capacity for fructan degradation (NES = −1.85, FDR = 0.019). The top sparse features on comp1 were the carbohydrate-binding module CBM97 and the glycoside hydrolases GH143, GH137, and GH138, all associated with pectin or hemicellulose substrates (Figure 3**I**).

Corroborating these multivariate findings, univariate ANCOVA models were fitted for 171 species, 1,671 KO terms, and 476 CAZymes independently. No individual feature survived FDR < 0.1 in any layer, non-expectedly given the limited sample size. Nevertheless, the ranking of features was highly concordant between the two frameworks: Spearman correlations between univariate T-statistics and PLS-DA component 1 loadings were r = 0.59 (species), r = 0.65 (KO), and r = 0.79 (CAZymes). Concordance at the pathway-enrichment level was high for species (r_NES_ = 0.73) and CAZymes (r_NES_ = 0.80), and moderate for KO pathways (r_NES_ = 0.39).

### 7.5. *Giardia* burden is reflected in community-wide bacterial composition

The analyses above characterize LBHF-associated shifts in microbiome composition. We next asked whether there are specific microbial configurations linked to *Giardia* burden. Since *Giardia* was resolved in almost all animals by day 45 (26/30), we conducted this analysis only using baseline samples.

Overall, no specific species abundance was significantly associated with *Giardia* abundance (FDR>0.05). When looking into taxonomic enrichments, we observed a total of 16 taxonomic levels related to *Giardia* concentrations. These included a negative relationship with Bacillota (NES = -1.91, FDR= 0.013) and Lachnospiraceae (NES = -1.79, FDR = 0.013), and positive enrichment of Fusobacteriaceae (NES = 1.97, FDR=0.013) and Bacteroidales (NES=1.76, FDR = 0.02). In addition, we identified a negative trend in which animals with higher *Giardia* have on average a lower alpha diversity (effect=-0.055, P=0.08).

We next assessed whether bacterial community composition could predict *Giardia* concentration. The microbial community showed modest predictive ability for *Giardia* load (XGBoost, LOOCV: Spearman’s *ρ* = 0.52, *P* = 0.002) (**Figure** 4**A**). SHAP-based feature attribution identified a restricted set of taxa as the principal drivers of model output (**Figure 4B**). Among these, *Turicibacter* sp002311155 emerged as the dominant contributor, with a mean absolute SHAP importance 12.4 standard deviations above the average taxon-level importance across the community. Additional influential taxa included *Allobaculum stercoricanis* (5.02 standard deviations above the mean), *Blautia* sp900556555 (4.53), *Megamonas* cluster34 (4.42), and CCUG_7971 cluster10 (3.63), whereas no other taxon exceeded two standard deviations above the community-wide mean importance. To examine directionality, we correlated taxon abundance with the corresponding taxon-specific SHAP values. Higher abundance of *Turicibacter* sp002311155, *Allobaculum stercoricanis*, and *Blautia* sp900556555 was associated with more negative SHAP values, indicating that these taxa tended to decrease predicted *Giardia* concentration. Conversely, higher abundance of *Megamonas* cluster34 and CCUG_7971 cluster10 was associated with more positive SHAP values, indicating that they tended to increase the predicted Giardia concentration (**Figure** 4**C**).

**Figure 4.**
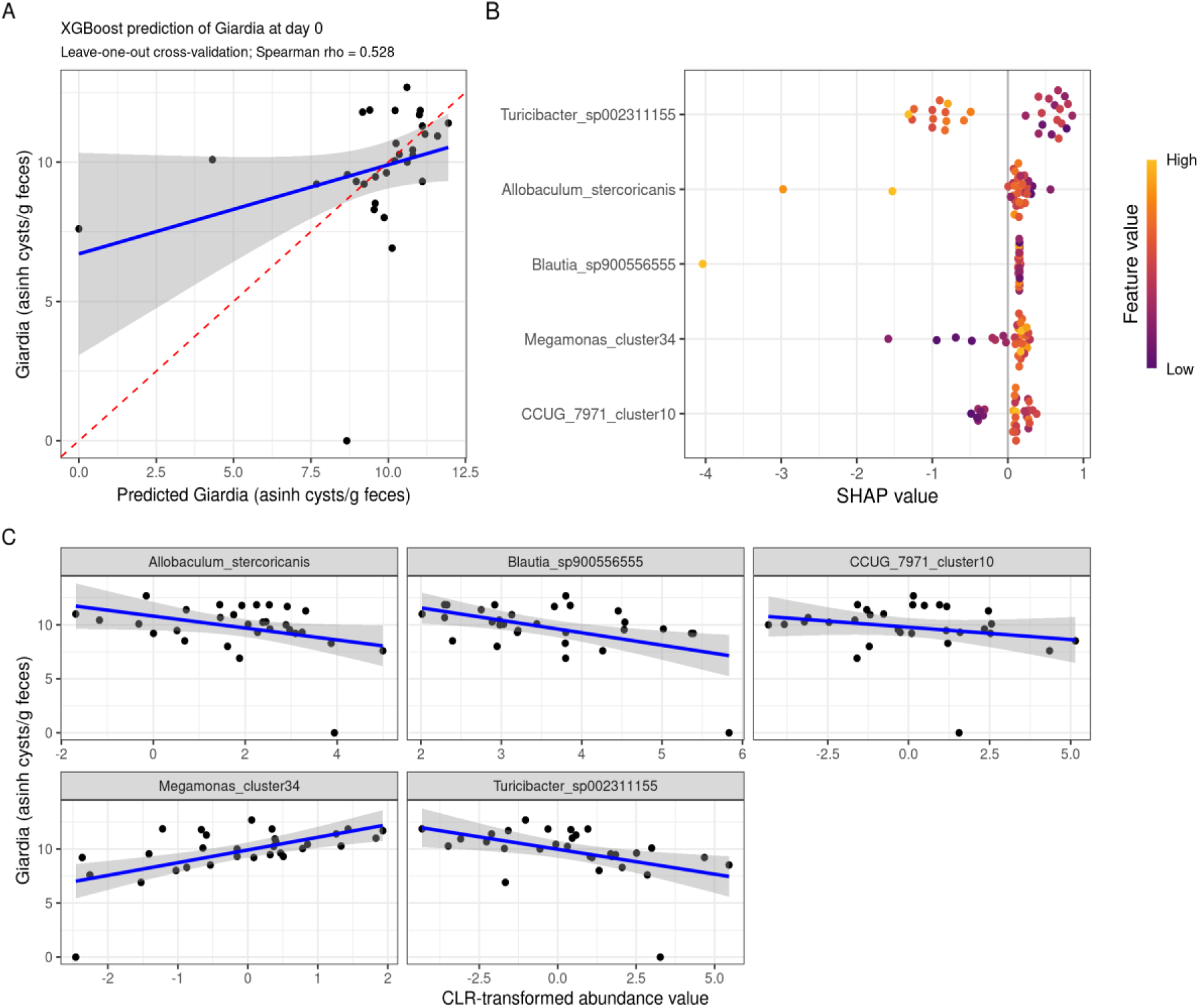
Giardia duodenalis concentration relates to gut microbial composition. (A) Prediction of Giardia concentration (asinh-transformed) using microbiome abundance at baseline (CLR-transformed). Predicted Giardia indicates the abundance predicted using an XGBoost model in a leave-one-out cross-validation framework per each of the test points. (B) SHAP-based feature attribution identifying the principal taxa driving the XGBoost model predictions. Only those taxa which absolute average importance over 2 standard deviations from the average taxon importance are shown. (C) Scatter plot shows the relationship between the abundance of the top species contributing to the model and Giardia.

## 8. Discussion

In this feeding study of juvenile dogs with confirmed natural *Giardia duodenalis* infection, daily supplementation with postbiotic LB fermentate HF (LBHF), adjunctively with metronidazole, accelerated parasite clearance compared to placebo. By D10, *Giardia* was cleared in all LBHF-treated dogs, whereas no dog in the placebo group receiving the same antiparasitic treatment had cleared the infection. Clearance in the LBHF group was sustained through D45 and was accompanied by improved stool consistency and lower fecal odor. Together, these observations support an adjuvant effect of LBHF under the conditions of this study, while not establishing the efficacy of LBHF as a stand-alone treatment.

Interpretation of the parasitological outcome should account for baseline differences in cyst shedding (numerically average lower cyst counts in the LBHF group) despite randomization. These differences were addressed analytically using individual baseline values as covariates. Importantly, individual dogs in the LBHF group with baseline *Giardia* cyst loads as high as in the placebo group also cleared the parasite by D10, whereas clearance was not observed in placebo-treated dogs at that time (Fig. 2).

The mechanism underlying accelerated clearance remains unresolved. *Giardia* control is strongly dependent on mucosal defenses, including *Giardia*-specific secretory IgA (Eckmann, 2003; Langford et al., 2002). In the present study, while no significant differences in total fecal sIgA were observed between groups, the LBHF-treated group tended to have lower levels, rather than higher. However, since we measured total sIgA and not anti-parasitic IgA, we cannot exclude nor confirm that LBHF had stimulated specific sIgA responses. The modest, non-significant decreases in total sIgA and fecal calprotectin could instead be compatible with reduced intestinal inflammatory activity in the LBHF group (Melcher et al., 2025), particularly in light of the improved fecal scores. Early sampling of *Giardia*-specific IgA, mucosal cytokines, epithelial barrier markers, and innate immune pathways will be needed to distinguish immune-mediated clearance from other mechanisms.

LBHF did not significantly change the canine dysbiosis index across the study, although it showed a trend for reduced dysbiosis. This index is derived from quantitative measurements of eight bacterial taxa and has been associated with disease severity in canine chronic inflammatory enteropathy. Its relevance to giardiasis is less certain: one recent study found no significant difference between symptomatic and asymptomatic infected dogs, while earlier work reported microbiome alterations associated with infection. This mixed literature is biologically notable because *Giardia* can impair nutrient absorption and increase delivery of unabsorbed substrates, particularly proteins and lipids, to the hindgut (Farthing, 1993), which would be expected to affect microbiome composition and activity.

Metronidazole usage has been related to gut dysbiosis development. Thus, we wondered whether the usage of LBHF might result in healthier microbial communities. Metagenomic profiling at baseline and D45 identified a coordinated LBHF-associated shift across taxonomy, functional gene repertoire, and carbohydrate-active enzymes. Relative increases in Bacteroidota, including *Prevotella*- and *Alloprevotella*-affiliated taxa, were accompanied by relative decreases in Bacilli, Campylobacterales, and Clostridia. Functional profiles indicated greater representation of pathways for degradation of plant-derived polysaccharides and lower representation of genes related to bacterial motility, biofilm formation, and Gram-positive cell-wall biosynthesis. Because microbiome sampling was performed only at baseline and D45, after all dogs had cleared *Giardia*, these data cannot determine whether the community shift caused, facilitated, or followed parasite clearance. Microbiome observations may nevertheless help contextualize the improvement in fecal quality and fecal odor. The suggested shift in GH families toward saccharolytic substrate utilization is consistent with the observed enrichment of *Prevotella*- and *Alloprevotella*-affiliated organisms and could plausibly reduce odor-generating products from proteolytic fermentation. The latter could also be related to the observed reduction in *Clostridia*. Reduced representation of flagellar assembly genes may also be relevant to intestinal inflammatory tone, although any link to stool quality remains speculative. Likewise, the observed negative association between *Allobaculum stercoricanis* abundance and *Giardia* burden should be considered hypothesis-generating because directionality cannot be established from the present sampling design.

Bile acid metabolism represents another plausible but unproven pathway for regulation of Giardia burden. *Turicibacter* has been reported to modulate bile acids and improve intestinal homeostasis, and in our study higher numbers were associated with lower Giardia counts. Secondly, although enzymatic activity is unlikely to remain highly active in a heat-treated postbiotic such as LBHF, Lactobacilli can express bile salt hydrolases (BSH), and deconjugated bile salts may be unfavorable to *Giardia.* However, fecal taxonomic and functional analyses did not indicate altered BSH representation. Nevertheless, an effect localized to mucosa-associated communities in the upper small intestine cannot be excluded because fecal metagenomics may not capture changes at the principal site of *Giardia* colonization. Therefore, it remains possible that components in LBHF may have affected small intestinal mucosal BSH production or may have interfered with enterohepatic bile salt circulation.

A more direct explanation is interference with parasite attachment or with the epithelial niche. Components of LBHF adhere to intestinal epithelial cells (Chauvière et al., 1992), and in vitro studies have reported inhibition of pathogen adhesion by LBHF cell bodies and metabolites (Coconnier et al., 1997, 2000; Liévin-Le Moal, 2016; Liévin-Le Moal et al., 2002, 2007). Related anti-pathogenic effects have been described in vivo (Liévin-Le Moal, 2016), including reduced *Helicobacter pylori* burden in a human trial with a postbiotic preparation with similar composition as LBHF (Canducci et al., 2000). Interestingly, in the present study, whereas none of the fecal samples contained *H. pylori*, other *Helicobacter* species were reduced with LBHF treatment as were KEGG pathways involved in *H. pylori* pathogenicity. LBHF was recently also found to reduce infectious diarrhea and mucosal pathology in pigs challenged with enterotoxigenic *Escherichia coli* (Duarte et al., 2024; Xu et al., 2022). Additionally, improved epithelial integrity associated with an observed increase in lipoarabinomannan biosynthesis activity could also make attachment sites less accessible to pathogens. These observations provide biological plausibility, but they do not demonstrate anti-adhesive activity against *Giardia*. Direct adhesion assays using *Giardia* trophozoites and epithelial models, followed by appropriately controlled in vivo studies, are needed to test this mechanism.

Despite limitations such as the baseline imbalance in *Giardia* shedding, our study nevertheless shows that LBHF can reduce Giardia burden and, importantly, reduce the breakdown in fecal scoring and malodor, which likely is a consequence of Giardiasis itself and dysbiotic effects of metronidazole. Larger, prospectively powered studies with balanced baseline burden, factorial treatment arms, earlier and repeated microbiome and mucosal sampling, and direct mechanistic assays would be needed to confirm the clinical effect and establish how LBHF accelerates clearance.

## 9. Conclusion

Adjunctive LBHF supplementation was associated with earlier clearance of natural *Giardia* infection and improved fecal quality and fecal odor in metronidazole-treated juvenile dogs. The study also identified a distinct day 45 microbiome configuration, but the available sampling cannot unequivocally establish whether these changes mediated parasite clearance. The findings justify confirmatory trials designed to separate treatment effects, resolve early mucosal and microbiome responses, and directly test anti-adhesive and immune mechanisms.

## 10. Acknowledgements

The authors gratefully acknowledge Maëlys Ferreira (Clinical Biology and Biochemistry) for her support and contributions to this study. We also thank all breeders who participated in the study for their cooperation and commitment.

## 11. Author contributions

E.E. – Review and editing, conceptualization, investigation, project administration, supervision

S. A-S. – Methodology, formal analysis, writing original draft, review and editing

B.T. – Methodology, formal analysis, review and editing

A.S.B. – Methodology, investigation

E.P. – Conceptualization, project administration

D.E. – data curation, formal analysis, investigation, methodology, project administration, resources, supervision, validation, writing – original draft, writing – review C editing

A.C. – data curation, formal analysis, investigation, methodology, project administration, resources, supervision, validation, writing – original draft, writing – review C editing

M.K. – Conceptualization, writing – review C editing

B. L. – data curation, investigation, methodology, writing – review C editing

P. D. – Conceptualization, data curation, formal analysis, investigation, methodology, validation, writing – review C editing

D. LR. – Conceptualization, data curation, formal analysis, investigation, methodology, validation, writing – review C editing

N. C-F. – Conceptualization, data curation, formal analysis, investigation, methodology, resources, supervision, validation, writing – review C editing

B. P. – Conceptualization, data curation, formal analysis, investigation, methodology, project administration, resources, supervision, validation, writing – review C editing

M.M. – Conceptualization, data curation, formal analysis, investigation, methodology, project administration, resources, supervision, validation, writing – original draft, writing – review C editing

## 13. Data availability

Raw sequencing data have been deposited in the NCBI Sequence Read Archive under BioProject accession PRJNA1516229 and will be made publicly available upon publication. https://dataview.ncbi.nlm.nih.gov/object/PRJNA1516229?reviewer=s5tta8pt2hs96csen809m8410l

